# Structure and function of the SIT1 proline transporter in complex with the COVID-19 receptor ACE2

**DOI:** 10.1101/2023.05.17.541173

**Authors:** Huanyu Z. Li, Ashley C.W. Pike, Gamma Chi, Jesper S. Hansen, Sarah G. Lee, Karin E.J. Rödström, Simon R. Bushell, David Speedman, Adam Evans, Dong Wang, Didi He, Leela Shrestha, Chady Nasrallah, Nicola A. Burgess-Brown, Timothy R. Dafforn, Elisabeth P. Carpenter, David B. Sauer

## Abstract

Proline is widely known as the only proteogenic amino acid with a secondary amine. In addition to its crucial role in protein structure, the secondary amino acid modulates neurotransmission and regulates the kinetics of signaling proteins. To understand the structural basis of proline import, we solved the structure of the proline transporter SIT1 in complex with the COVID-19 viral receptor ACE2 by cryo-electron microscopy. The structure of pipecolate-bound SIT1 reveals the specific sequence requirements for proline transport in the SLC6 family and how this protein excludes amino acids with extended side chains. By comparing apo and substrate-bound SIT1 states, we also identify the structural changes which link substrate release and opening of the cytoplasmic gate, and provide an explanation for how a missense mutation in the transporter causes iminoglycinuria.

## Background

Proline is the only amino acid incorporated into proteins that lacks a primary amine group. With its pyrrolidine ring, the amino acid’s restricted Ramachandran angles and hydrogen bonding capability have pronounced effects on polypeptide secondary structure ^1^. Consequently, the residue is usually found at the ends of alpha helices and at bends in helices where it disrupts the hydrogen bonding pattern ^2^, while proline and its derivate hydroxyproline are overrepresented in Polyproline-II helices and the collagen triple helix ^3, 4^. Within a protein, proline’s unique cis/trans isomer energetics and isomerization kinetics are central to kinetic switches in signaling proteins ^5, 6^, and protein folding ^7^. Physiologically, the amino acid also acts as a weak agonist of glycine and ionotropic glutamate receptors ^8, 9^, and hyperprolinemia is associated with autism spectrum disorder, intellectual disability, and psychosis spectrum disorders ^10^.

Several plasma membrane transporters import proline into the cell, including the Sodium/imino-acid transporter 1 (SIT1) encoded by the SLC6A20 gene. SIT1 was first identified as a proline transporter in the kidney ^11–13^. Consequently, polymorphisms in SLC6A20 lead to iminoglycinuria ^14^, and are correlated with altered concentrations of secondary and tertiary amine metabolites in plasma and urine ^15, 16^. Neurologically, SIT1 regulates proline concentrations to modulate the activity of glycine and NMDA-type glutamate receptors in mice ^9^, and the absence of neurons in the colon, causing Hirschsprung’s disease, is associated with SLC6A20 polymorphisms ^17–19^. In the eye, SIT1 expression is a signature of the retinal pigment epithelium and drives the proline-preferring metabolism of these cells ^20–22^. Accordingly, gene variants are correlated with both retinal and macular thickness, and degenerative macular disease ^21, 23, 24^. Finally, SIT1 traffics to the plasma membrane in a complex with ACE2 ^25^, the SARS-CoV2 receptor. SIT1 overexpression can prevent ACE2 trafficking to the plasma membrane ^26^ and polymorphisms in the transporter gene are associated with clinical outcomes of SARS-CoV2 infection ^27–29^

SIT1 belongs to the SLC6 gene family of amino acid and amine transporters ^30^, and the larger Neurotransmitter Sodium Symporter (NSS) superfamily. The structure, selectivity, and transport for SLC6 and NSS transporters has been revolutionized by structural studies of the prokaryotic amino acid transporters LeuT and MhsT ^31–36^, structurally homologous bacterial transporters Mhp1 and vSGLT ^37, 38^, and several eukaryotic SLC6 transporters ^39–43^. Central to NSS-mediated substrate transport is the

LeuT protein fold, a compact domain composed of 10 transmembrane helices ^31^. Within this structure, substrates and co-transported ions bind at sites created by breaks in TM1 and TM6, and differences in sequence within and near this region determine the proteins’ substrate selectivity ^35, 36^. Starting in an outward-open apo state, substrate and ion binding induce closure of the extracellular gate, through the movement of TM1b and TM6a and residues lining those helices which block access to the binding site ^31, 34^. From this occluded state, the cytoplasmic gate subsequently opens through tilting of TM1a into the plane of the bilayer ^34, 44^. Coupled to the tilting of TM1a are movements on the cytoplasmic face of the protein, particularly in gating helix TM5 with its highly conserved GX_N_P motif ^32^.

SIT1 and PROT (SLC6A7) are unique among the SLC6 family in preferring amino acids with a secondary amine ^11–13, 45^, though other SLC6 transporters can transport both primary and secondary amino acids ^30^. While the mechanism of substrate selectivity within the SLC6 family is of great interest, the proline transporters are relatively understudied. Furthermore, low sequence similarity limits useful comparison of SIT1 to the prokaryotic proline transporter PutP despite similar substrate selectivity profiles ^46^ (Supplementary Fig. 1a). Therefore, while homology models based on the prokaryotic LeuT structure have been used to probe SIT1’s ion binding ^47^, the mechanism for its selective transport of secondary amino acids remains unclear.

In this study, we probe SIT1’s selectivity and transport mechanism with a combination of thermostabilization-based SIT1 binding assays and cryo-electron microscopy (cryo-EM) based structural studies of the ACE2-SIT1 complex. From these results, we propose a structural model for SIT1’s preference for secondary amino acids and the conformational changes underlying amino acid release.

## Results

### Structure of ACE2-SIT1 complex

After overexpressing and purifying SIT1 (Supplementary Fig. 1b, c), we first validated substrate binding of SIT1 in detergent (Fig. 1a, Supplementary Fig. 1d). In agreement with the transporter’s *in vivo* selectivity ^13^, proline and pipecolate increased the protein melting temperature (T_M_) by 3 °C and 6 °C, respectively, while glycine and sarcosine had no apparent effect. Aiming to examine the structural interactions of SIT1 and substrate, the small size of the transporter-amino acid complex presented a challenge for single-particle cryo-EM. To increase the particle mass, we expressed and purified SIT1 in complex with its trafficking chaperone ACE2 (Supplementary Fig. 1e, f) ^25^, a strategy also used in determining the structure of apo-SIT1 and the neutral amino acid transporter B^0^AT1 ^26, 39–41^.

**Figure 1.**
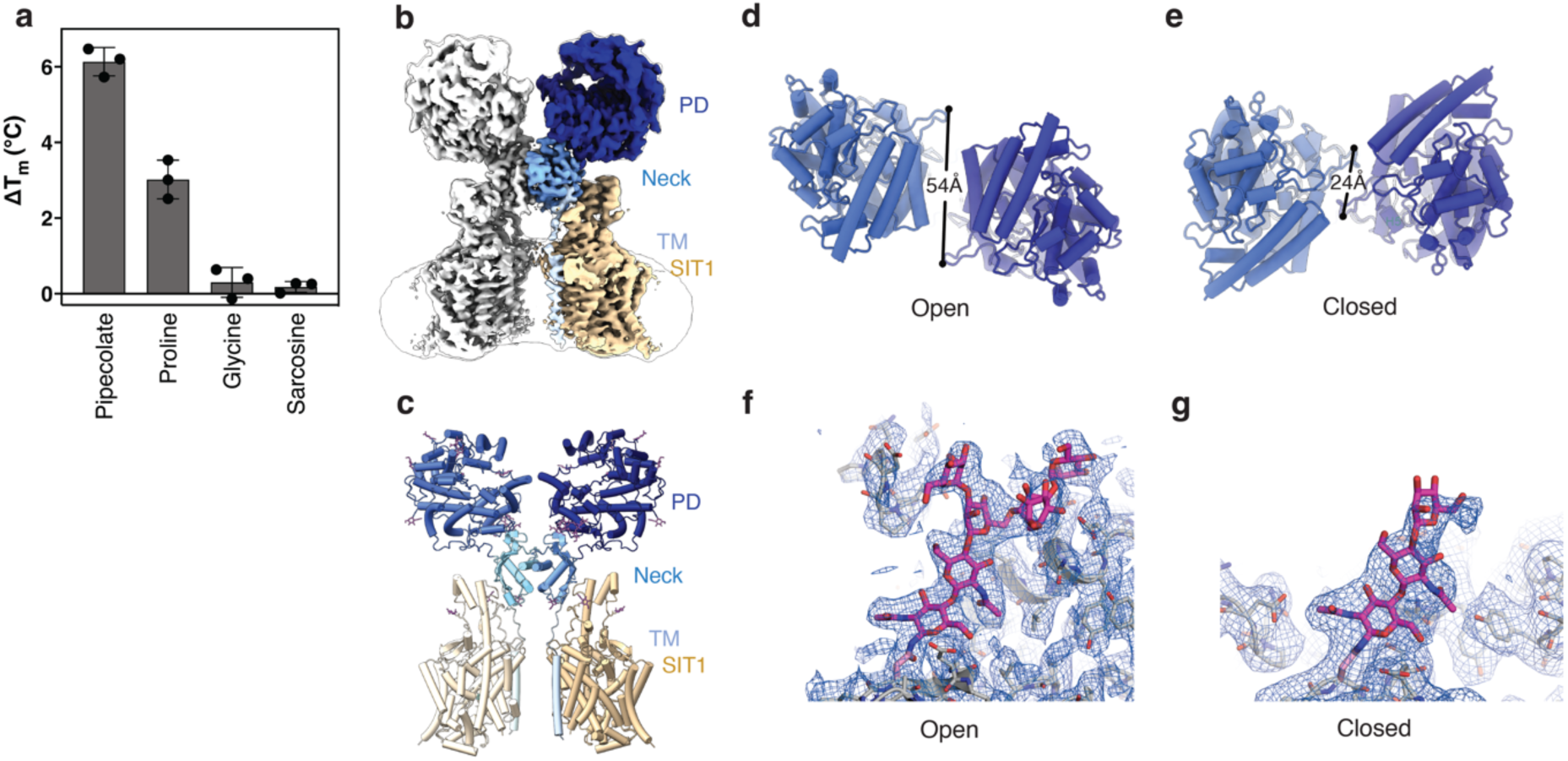
Structure of the ACE2-SIT1 complex determined in the presence of pipecolate by cryo-EM. (**a**) Change in melting temperature of SIT1 by amino acids. (**b**) Cryo-EM map of the ACE2-SIT1 complex determined in the presence of pipecolate. The ACE2 peptidase, neck, and TM domains are colored blue, cyan, and light blue, respectively. SIT1 is colored in wheat. Overlayed semi-transparent is the same map, low-pass filtered, showing the entire particle including the detergent micelle. (**c**) Protein structure of the ACE2-SIT1 complex, viewed from the plane of the membrane. Glycans are shown as purple sticks. ACE2’s peptidase domain in (**d**) open and (**e**) closed conformations. The glycan chain at Asn690, viewed from the membrane surface, interacting with the (**f**) open and (**g**) closed conformations of the peptidase domain with Coulombic potential maps shown as mesh.

Single-particle cryo-EM analysis ultimately yielded a nominal 3.24 Å map of the ACE2-SIT1 complex, determined in the presence of pipecolate (Fig. 1b, Supplementary Fig. 2a). This map was sufficiently detailed to model residues 10-582 of SIT1 and 21-768 of ACE2. SIT1 adopts the classic LeuT-fold expected for this family of amino acid transporters ^26, 47^, while ACE2 is composed of peptidase (PD) domain and collectrin-like domain with transmembrane and neck regions ^39^. As with the homologous ACE2-B^0^AT1 complex, dimerization of ACE2 is mediated primarily by its neck domain, while the ACE2 and transporter subunits interact via three distinct sets of contacts (Fig. 1c, Supplementary Fig. 3a-c). Within the membrane, the transmembrane helix of ACE2 makes extensive van der Waals contacts with TM3 and TM4 of SIT1. On the extracellular side of the membrane ECL2 of SIT1 hydrogen bonds with the extended region between the neck and TM domain of ACE2. Finally, the C-terminal portion of TM7 and loop prior to EH5 from SIT1 hydrogen bond with the collectrin-like domain of ACE2. The transporter is N-glycosylated at Asn131 and Asn357. ACE2 is more heavily decorated ^48^, with visible N-linked glycans at Asn90, Asn103, Asn322, Asn432, Asn546, Asn690, and an O-glycosylation at Thr730 (Fig. 1c, Supplementary Fig. 3d). Most of these glycan chains do not significantly interact with the protein, and consequently only a single sugar is resolvable. However, a branched N-linked glycan at ACE2’s Asn690 extensively hydrogen bonds with the peptidase domain.

While processing and classifying particles from the ACE2-SIT1 sample, obvious structural and compositional heterogeneity was apparent within the dataset (Supplementary Fig. 2a). Unlike previous ACE2 complex structures, this ACE2-SIT1 dataset yielded maps with 2:2 and 2:1 stoichiometry of ACE2 to transporter, with 3-fold more particles in the larger complex. This varying stoichiometry is not due to insufficient SIT1 as un-bound transporter was apparent during purification (Supplementary Fig. 1f). We hypothesize this smaller complex may be a consequence of low affinity between the SIT1 and ACE2 leading to varying stoichiometry not resolvable by preparative size exclusion chromatography. Alternatively, this structural heterogeneity may be due to the denaturation of a single transporter subunit at the air-water interface ^49^.

Significant structural heterogeneity in the ACE2’s peptidase domain was present within the ACE2-SIT1 complex. Further classifying the 2:2 ACE2-SIT1 complex data, there are two distinct conformations based on the relative domain orientations within the ACE2 dimer. This movement between open and closed conformations occurs as a 30° rigid-body rotation around a hinge at residues 612-617 in the loop between peptidase and collectrin-like domains (Supplementary Fig. 3e-g). As with the ACE2-B^0^AT1 structure ^39^, a second dimer interface is formed by Gln139 and Gln175 within the peptidase domain in ACE2’s closed conformation (Fig. 1d, e). Further, the branched N-glycan chain at ACE2’s Asn690 is rigid and well-ordered in the open conformation, interacting with the peptidase domain’s H6 and beta-hairpin between H4 and H5 (Fig. 1f). These structural features of the PD move in the closed conformation, breaking the protein-glycan interactions. Accordingly, the glycan density is weaker in this conformation as the glycan chain becomes more mobile (Fig. 1g). This suggests that the Asn690 glycan chain acts as a latch on the PD domain, stabilizing the open conformation and thereby regulating its change between conformations.

### Substrates binding within SIT1

While the 2:2 ACE2-SIT1 complex with the peptidase domain in the closed conformation yielded the highest-resolution map for the entire complex, the map is best in ACE2’s neck region and less detailed in the transporter (Supplementary Fig. 2a). We therefore combined particles from closed and open conformations, symmetry expanded over the transmembrane region of ACE2 and SIT1, and performed a further round of 3D classification and local refinement. This strategy produced a new 3.49 Å map of the transmembrane domain. Map interpretability improved for the peptide backbone and side chains (Supplementary Fig. 2b-d), enabling more accurate placement for residues 10-582 of SIT1 and 740-768 of ACE2’s TMD.

SIT1 is in an occluded conformation with the active site inaccessible to both the cytoplasm and extracellular space. The extracellular gate is closed with Tyr102 and Phe250 blocking access to the substrate binding site. The extracellular path to the substrate site is further obstructed by Tyr33 and Asn461, which are hydrogen bonded, and potential water-mediated hydrogen bonds between Arg30, Tyr38, Asn243, and Asp462. On the opposite side of the membrane, the cytoplasmic gate formed by TM1a is stabilized primarily through van der Waals interactions with TM6b and TM7. This gate is further held closed by two hydrogen bond networks linking TM1a’s Ser11 with Asn270 and His275 on TM6, and Tyr21 with Gly253 and Ser258 on TM7.

Within the SIT1 binding site are two non-protein densities, identified as pipecolate and chloride based on size, local chemistry, and similarity to structures of other LeuT-fold transporters (Fig. 2a, d). SIT1 engages the distinct chemical moieties of pipecolate through three regions of the binding site. The pipecolate’s amino group is surrounded by the carbonyls of Tyr21 and Ala22 of TM1 and Phe250 and Ser251 of TM6 (Fig. 2b), with Ala22 and Ser251 best placed for hydrogen bonds. The substrate’s carboxy moiety interacts with the amide nitrogens of Gly24, Leu25, and Gly26, and the side chain hydroxyl from Tyr102 of TM3 (Fig. 2b). Finally, the substrate’s piperidine ring is coordinated by van der Waals contact from Tyr21 of TM1, Leu98 of TM3, Phe250, Gly253, and Phe256 of TM6, and Asn410 of TM8 (Fig. 2c).

**Figure 2.**
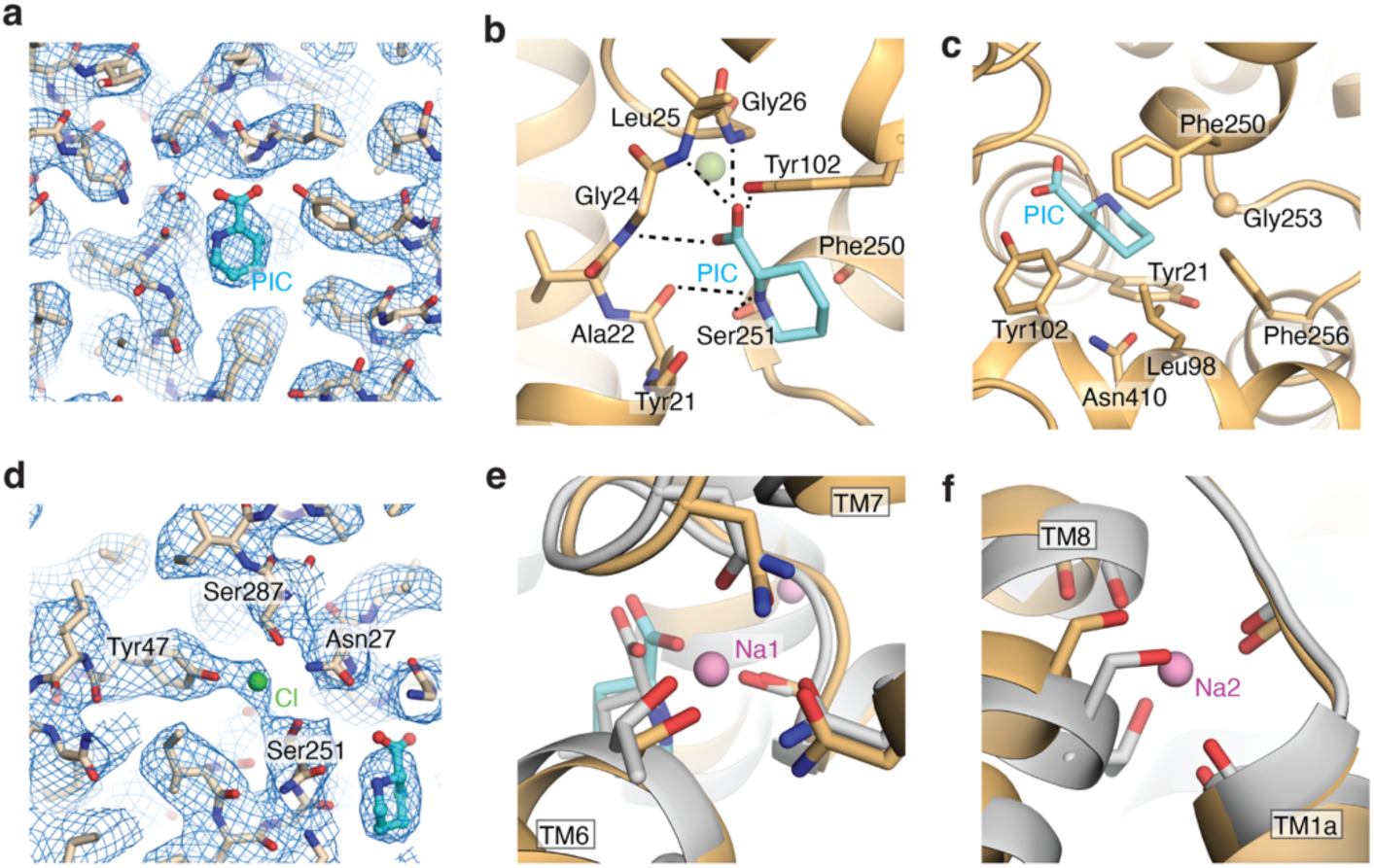
Amino acid, chloride, and sodium binding sites of SIT1. (**a**) Amino acid binding site of SIT1. Coulombic potential map shown as mesh, pipecolate shown as cyan sticks. SIT1’s coordination of the pipecolate’s (**b**) amine and carboxylate groups and (**c**) piperidine ring. (**d**) Chloride binding site of SIT1. Coulombic potential map shown as mesh. Overlay of the (**e**) Na1 and (**f**) Na2 binding sites for SIT1 and LeuT. SIT1 and LeuT are shown in wheat and grey, respectively. Sodium ions from the LeuT structure (2A65) are shown as purple spheres.

SIT1’s binding of pipecolate is very similar to the amino acid coordination by the S1 binding site of LeuT and MhsT ^31, 32^ (Supplementary Fig. 4a, d, g). In contrast, the pose of leucine in the related B^0^AT1’s structure is significantly different from that observed here, or in LeuT and MhsT (Supplementary Fig. 4b, e). However, the leucine-bound ACE2-B^0^AT1-focused cryo-EM map (EMD-30043) contains ambiguous densities for the substrate. Refitting the leucine and binding site residues in B^0^AT1 to positions more consistent with homologous transporters (Supplementary Fig. 4c, f) yielded an improved substrate FSC-Q ^50^. Therefore, we used this re-refined model of ACE2-B^0^AT1 bound to leucine for all subsequent comparisons.

As a sodium and chloride coupled co-transporter, SIT1 has obvious binding sites for two sodium and one chloride ion. The density for the Cl^-^ is clear in the experimental Columbic potential map (Fig. 2d). Within SIT1, the anion is coordinated by the side chains of Asn27 of TM1, Tyr47 of TM2, Gln247 and Ser251 of TM6, and Ser287 of TM7 in a mode very similar to hSERT and the engineered, chloride-dependent LeuT ^43, 51^. Notably, there is no apparent density within SIT1 for sodium ions at the expected Na1 and Na2 sites, despite an experimental concentration 7-fold greater than the cation’s K_M_ ^47^. However, sodium density is often weak or absent in cryo-EM maps at similar or better resolution ^52–54^. Nevertheless, the coordinating moieties from SIT1 and substrate are oriented similarly to the sodium-bound state of LeuT (Fig. 2e, f). Furthermore, the valence at the Na1 (v_Na_ = 2.5) and Na2 (v_Na_ = 0.41) sites indicate reasonable coordination for sodium ions ^55^. Therefore, we propose this structure captures the sodium, chloride, and substrate-bound inward-facing occluded state of SIT1’s reaction cycle.

### Proline binding in the SLC6 family

To understand the capability and preference for SLC6 transporters to import proline, we compared the binding site interactions of SIT1 to the structures and sequences of the related transporters. As previously noted, Gly253 packs immediately against the piperidine ring of pipecolate (Fig. 3a), and other SLC6 transporters of proline all possess a glycine at the equivalent position (Fig. 3c). In contrast, nearly all of the remaining amino acid transporting SLC6s have a serine at this position, which make hydrogen bonds with the substrates’ primary amine in B^0^AT1, LeuT, MhsT (Fig. 3b) ^31, 32^. Such a serine in SIT1 would sterically clash with proline in the binding site (Supplementary Fig. 4h). Therefore, we propose that SLC6 proteins require a small side chain at this position for secondary amino acid transport.

**Figure 3.**
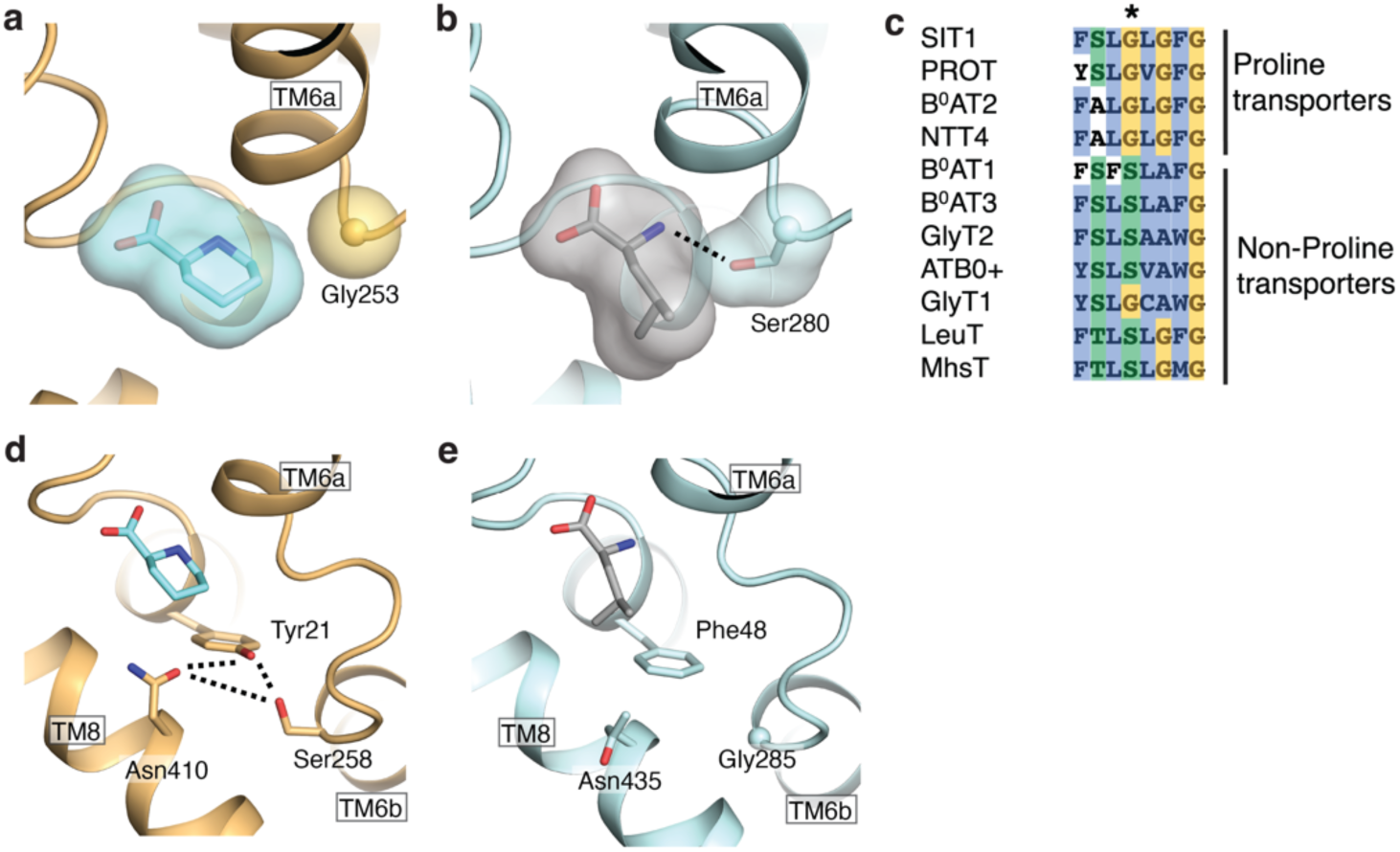
Binding site differences between proline selective and non-selective SLC6 amino acid transporters. Binding site for (**a**) SIT1 and (**b**) the refit B^0^AT1, colored in wheat and light blue, respectively. Substrate pipecolate and leucine are shown in cyan and grey, respectively. (**c**) Sequence alignment of SLC6 amino acid transporters. The position of SIT1’s Gly253 is indicated. Hydrogen bond network of the pipecolate coordinating residues Tyr21 and Asn410 (**e**) in SIT1 and (**f**) equivalent region in B^0^AT1.

While SIT1’s Gly253 explains a necessary component of secondary amine transport, this mechanism does not explain SIT1’s exclusion of amino acids with extended side chains ^13^. Examining the substrate binding site, we noted pipecolate is contacted by the side chains of Tyr21 and Asn410, which are hydrogen bonded to each other and Ser258 (Fig. 3d). This network restrains the position of Asn410 such that it would clash with amino acid substrates possessing extended side chains. This immediately suggests a mechanism for SIT1’s selectivity, where Asn410 is a steric block to exclude substrate amino acids with extended sidechains. Consistent with this model, the neutral amino acid transporting B^0^AT1 lacks this hydrogen bond, with phenylalanine at the position equivalent to SIT1’s Tyr21. Therefore, B^0^AT1’s equivalent asparagine, Asn435, is free to move and thereby accommodate the substrate’s extended side chain, as seen in protein’s leucine-bound structure (Fig. 3e, Supplementary Fig. 4h). Furthermore, PROT also appears to use a similar steric block to exclude extended side-chain substrates. Rather than the hydrogen bond of SIT1, PROT appears to use the bulk of a phenylalanine at the equivalent position to Asn410 to prevent access to the side chain pocket (Supplementary Fig. 4i).

### Opening of SIT1’s cytoplasmic gate in the absence of substrate

Having captured the inward-facing occluded state of SIT1’s reaction cycle, we next set out to characterize the protein’s conformational changes upon substrate release. We therefore determined the ACE2-SIT1 structure, with an open-conformation peptidase domain, in the presence of the glycine to a resolution of 3.46 Å overall and 3.76 Å for the transporter alone (Supplementary Fig. 5, Supplementary Fig. 6a). The maps were sufficiently detailed to model and refine residues 11-582 of SIT1 and 20-768 of ACE2 (Supplementary Fig. 6b-d) This SIT1 structure agrees well with the recently published ACE2-SIT1 structures in complex with receptor binding domains from COVID-19 and determined in amino acid-free buffers (RMSD = 0.765-0.940).

Without a secondary amino acid substrate, SIT1 has undergone structural rearrangements that open the transporter’s cytoplasmic gate (Fig. 4a, b). The greatest movement is a rigid body 17° tilt of TM1a (Fig. 4c). This movement breaks most of TM1a’s closed-conformation hydrophobic interactions with TM6b and TM7, and the hydrogen bonds of Ser11 (Fig. 4d). The absence of density for Tyr21 indicates it is mobile and therefore no longer hydrogen bonded to Ser258 on TM6b. Rather, in this inward-open conformation TM1a forms exclusively van der Waals contacts with TM5 and TM7 (Fig. 4e). This agrees with the weaker TM1a interactions in other LeuT-fold transporters upon substrate release which increase the dynamics of this helix ^56^.

**Figure 4.**
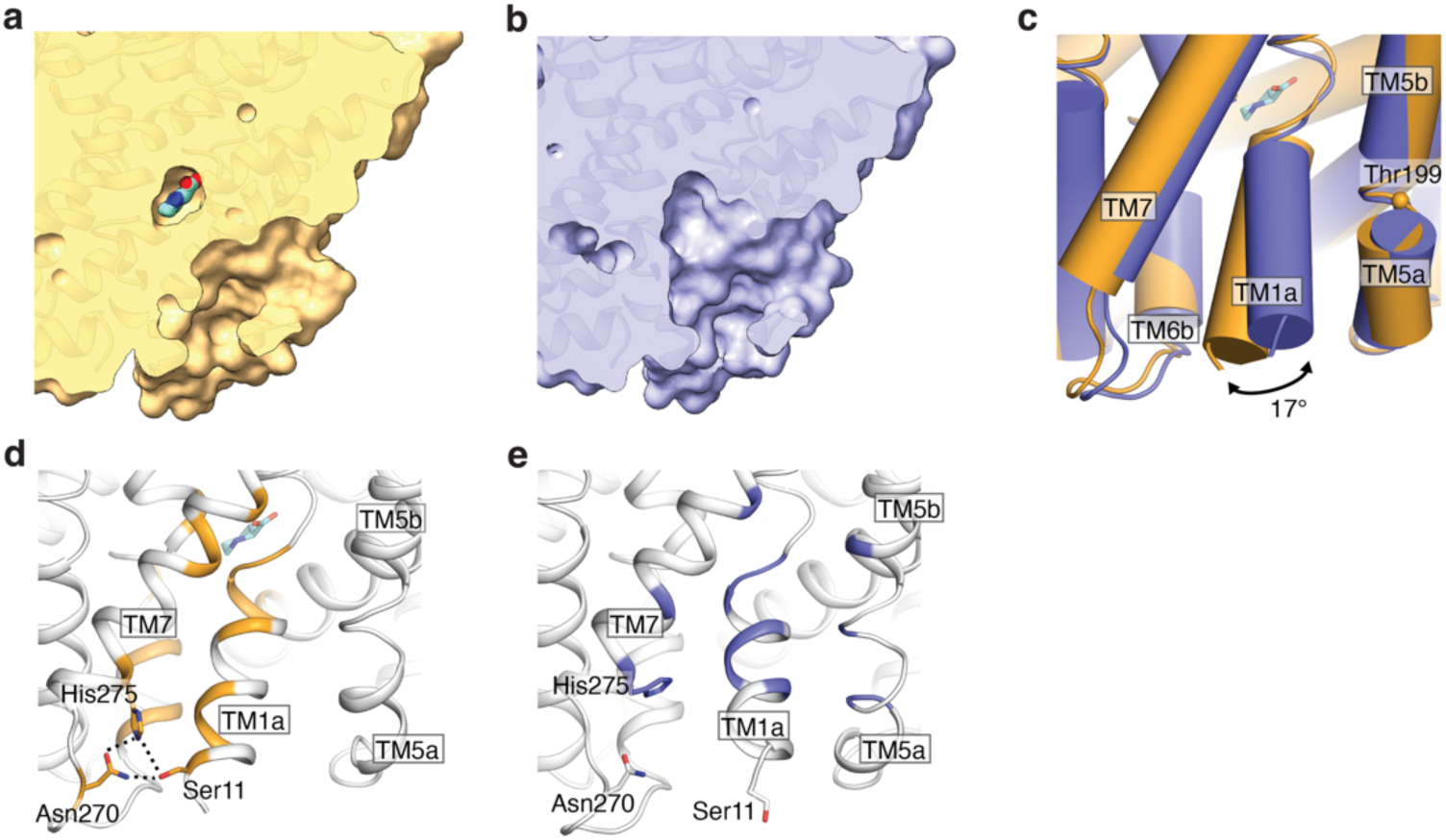
Cryo-EM structure of the ACE2-SIT1 complex determined in the inward-facing apo state. Cross section of the ACE2-SIT1 transmembrane domain in (**a**) pipecolate-bound and (**b**) apo state, determined in the presence of glycine. (**c**) Overlay of SIT1 in the pipecolate-bound and apo states. (**d**) The structure of pipecolate-bound SIT1, with residues within 3.9 Å of TM1a highlighted in wheat. (**e**) The structure of apo SIT1, with residues within 3.9 Å of TM1a shown in purple. TM1a interacting residues for SIT1 in the (**d**) pipecolate-bound (**e**) apo state, determined in the presence of glycine. Residues within 3.9 Å of TM1a are highlighted in wheat and purple, respectively.

### Mechanism of the iminoglycinuria mutation

The open conformation of SIT1’s cytoplasmic gate is stabilized by TM1a’s interactions with TM5, which has moved laterally within the membrane (Fig. 4c, e). This mobile portion of TM5 corresponds with the conserved GX_N_P motif essential to opening the cytoplasmic gate of LeuT-fold transporters ^32^. This immediately suggested a mechanistic explanation for the mutation in TM5, T199M, implicated in iminoglycinuria (Fig. 4c) ^14^. We expect the larger methionine side chain in the mutation to interfere with the packing and dynamics of TM5. This would then alter SIT1’s energetics for opening the cytoplasmic gate and thereby reduce the proline transport rate. Supporting this hypothesis, the T199M mutation produces an eight-fold reduction in SIT1’s proline transport Vmax ^14^. This mechanism is also consistent with the partial rescue of SIT1’s iminoglycinuria mutant by the secondary mutation M401T ^47^. This mutation has been previously proposed to restore activity to the SIT1 T199M mutant by reestablishing a TM8-mediated link between TM5 and the amino acid and sodium sites. However, in our structures there is no change in TM8 upon substrate release. Rather, we propose the M401T mutation, in the background of T199M, allows TM5 to properly pack against TM8 and thereby restores the energetics of opening SIT1’s cytoplasmic gate.

### Structural coupling between TM1a and the substrate binding site

Within the SIT1 map determined in glycine, there is no apparent density for amino acid or chloride (Supplementary Fig. 6e, f). Rather, structural changes in this region appear to link the cytoplasmic gate to the sodium, chloride, and substrate sites. Most pronounced, the unwound region of TM1 has repositioned, distorting the hydrogen bond donors and acceptors which previously coordinated the substrate’s amine and carboxy groups (Fig. 5a). The side chain of Asn27 has shifted to partially occupy the Na1 site, also preventing its coordination of chloride (Fig. 5b, c). The tilting of TM1a has shifted the carbonyls of Ser20, Ala22, and Val23 away from both sodium sites (Fig. 5c, d). These structural distortions reduce the valence at the Na1 (v_Na_ = 0.26) and Na2 (v_Na_ = 0.08) sites, suggesting that both have lower affinities for sodium. The absence of Na1 may also alter the dynamics of Ser251 which coordinates both that cation and chloride, and an analogous link between substrate, sodium, and chloride binding has been proposed for an engineered LeuT ^51^.

**Figure 5.**
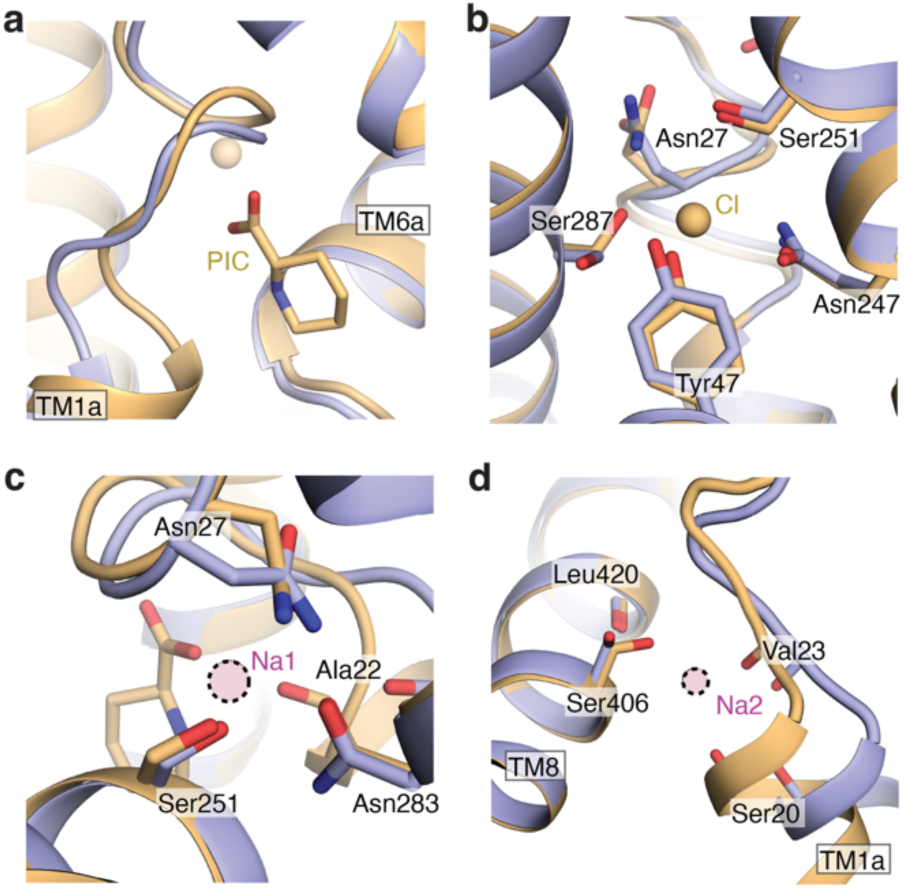
Substrate and ion binding sites of apo SIT1. (**a**) Overlay of SIT1’s pipecolate coordinating residues for the bound and apo structures. Pipecolate-bound and apo SIT1 are colored in wheat and purple, respectively. (**b**) Overlay of the pipecolate-bound and apo SIT1 chloride binding site. Overlay of pipecolate-bound and apo SIT1 for the (**d**) Na1 and (**e**) Na2 binding sites. Sodium positions are docked from sodium-bound inward-facing LeuT structure, and indicated by purple spheres with dotted edges.

**Table 1.**
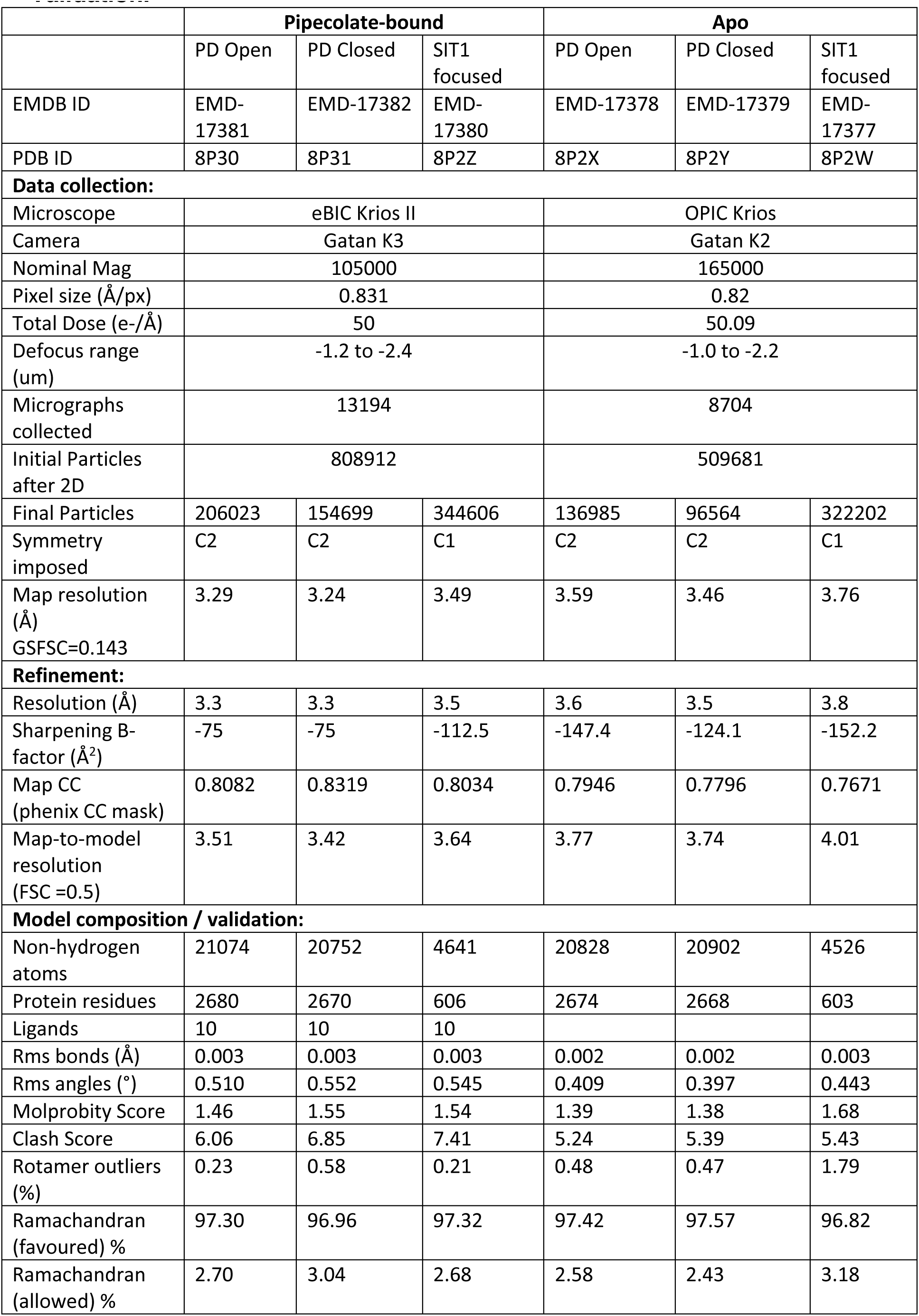
CryoEM Data Collection, Processing, Refinement and Model Validation.

## Discussion

Here we identify the structural features which determine the SLC6 proteins’ capability and preference to transport secondary amino acids, and the structural changes upon substrate release. Specifically, steric blocks mediated by a conserved glycine and asparagine explain SIT1’s selectivity for secondary amino acids. Further comparing SIT1’s apo and substrate-bound structures, we noted the release of substrate, sodium, and chloride, and the opening of the cytoplasmic gate, are synchronized through modest changes to protein structure in the binding site.

While excluding amino acids with extended side chains, SIT1’s steric block by Asn410 would allow binding and transport of amino acids with small side chains. This is consistent with the protein’s transport of sarcosine ^12, 13^, and the association of SLC6A20 gene variants with the concentration of dimethylalanine in urine ^15^. While this steric block also agrees with glycine transport by SIT1, there are conflicting reports for the transporter’s import of that amino acid ^9, 12, 13^. Our biochemical binding results were ambiguous, as both glycine and the *bona fide* SIT1 substrate sarcosine did not alter the transporter’s melting temperature (Fig. 1a, Supplementary Fig. 1d). This demonstrates the transporter’s interactions with small amino acids are insufficient to alter apparent stability, and thermostabilization cannot be used to exclude particular amino acids as substrates. However, our structural results do not support the transport of glycine. SIT1’s binding site adopts an inward-open apo state in the presence of that amino acid (Fig. 5b, c), despite being sterically accommodated when docked into the pipecolate-bound SIT1 structure. Notably, when docking sarcosine into the pipecolate-bound structure, its methyl group makes van der Waals contact with Tyr21, while bound glycine would lack this interaction (Fig. 2c, 3d).

The SIT1-pipecolate complex structure also suggests an explanation for the transporter’s preference for that amino acid over the smaller proline ^11^. Within the substrate binding site, Tyr21’s aromatic ring makes a van der Waals contact with the pipecolate (Fig. 3d), and the position of this tyrosine is restrained by the hydrogen bonds with Ser258 and Asn410. Therefore, we hypothesize this positional restraint prevents Tyr21 from making effective van der Waals contacts with smaller proline, explaining SIT1’s relative affinity. In contrast, PROT lacks the hydrogen bonding side chains on TM6 and TM8, equivalent to SIT1’s Ser258 and Asn410 (Supplementary Fig. 4i). Therefore, PROT’s Tyr53 can reposition for van der Waals contacts with smaller cyclic amino acids. Accordingly, PROT has a 2-fold tighter affinity for proline over pipecolate ^57^.

Finally, our structure and sequence provide hints for SIT1’s transport for N-Methyl-L-proline ^13^, while PROT cannot ^57^. This permissivity of SIT1 to N-methylated cyclic amino acids is also seen in GWAS analysis of circulating metabolites, where SLC6A20 gene variants are associated with the plasma concentrations of the tigonelline and N-methylpipecolate ^16^. Notably, within the SIT1-pipecolate structure the substrate’s amine nitrogen is engaged by Ala22 (Fig. 2a), and docking N-methylpipecolate into the structure produces a clash with that residue. We speculate the unwound region of TM1 in SIT1 can repack around the methyl groups of tertiary amine substrates, enabled by the small alanine side chain. However, the larger equivalent cysteine of PROT may block this structural accommodation, thereby explaining the proteins’ difference in selectivity for N-methylated cyclic amino acids.

## Methods

### Sequence alignment and phylogenomic analysis

SLC6 family protein sequences, and related bacterial homologs, were aligned in Promals3D ^58^. Phylogenetic distances were calculated using FastTree 2 with default settings ^59^, and rooted using NCC1 as the outgroup ^60^.

### Cloning

The full-length, codon-optimized sequence of human ACE2 was cloned into pHTBV (kindly provided by Prof. Frederick Boyce, Harvard) with N-terminal FLAG tag. The SIT1 sequence was cloned into pHTBV with C-terminal twin-Strep and 10-His tags.

Baculoviruses for each construct were generated using standard methods ^61^. Baculoviral DNA from transformed DH10Bac was used to transfect Sf9 cells to produce baculovirus particles, which were then amplified with Sf9 cells grown in Sf-900 II medium (Thermo Fisher Scientific) supplemented with 2% fetal bovine serum. Cells were incubated in an orbital shaker for 72 h at 27°C. Cultures were centrifuged at 900 g for 10 min to harvest the supernatants containing the viruses.

### SIT1 expression and purification

Expi293F GnTI− cells in Freestyle 293 Expression Medium (Thermo Fisher Scientific) were transduced with the SIT1 P3 baculovirus (3% v/v) in the presence of 5 mM sodium butyrate. Cells were grown in a humidity-controlled orbital shaker for 72 h at 30°C with 8% CO_2_ before being harvested by centrifugation at 900 g for 15 min, washed with phosphate-buffered saline, and flash-frozen in liquid nitrogen. Cell pellets were stored at −80°C until further use.

Cell pellets expressing full-length SIT1 were resuspended in a lysis buffer of 50 mM HEPES pH 7.5, 300 mM NaCl, 1.5% glycol-diosgenin (GDN, Anatrace), and cOmplete EDTA-free Protease Inhibitor Cocktail (Roche). Lysate was loaded on TALON resin (Takara Bio) gravity flow column, washed with column buffer (50 mM HEPES pH 7.5, 300 mM NaCl, 0.02% w/v GDN) supplemented with 1 mM ATP and 10 mM MgCl_2_, and eluted in column buffer with 300 mM imidazole. The eluent was immediately loaded on Strep-Tactin XT resin (IBA) gravity flow column, washed with column buffer supplemented with 1 mM ATP and 10 mM MgCl_2_, and eluted in column buffer with 50 mM D-biotin. The protein was further purified by size exclusion chromatography using a Superdex 200 Increase (10/300) GL column pre-equilibrated with SEC buffer (20 mM HEPES pH 7.5, 150 mM NaCl, 0.02% w/v GDN).

### Thermostabilization measurements

Purified SIT1 was diluted to 0.4 mg/mL in SEC buffer. Amino acids at 0.5 mM were added to the protein and incubated on ice for 1 hr, then Plex nanoDSF Grade High Sensitivity Capillaries (NanoTemper) were filled with 10-µl protein sample. Melting curves were determined using Prometheus NT.48 by monitoring the intrinsic fluorescence at 350nm relative to 330nm during a temperature ramp (1°C/min increase) from 20 to 95°C. The melting temperature was determined from measurements of three biological replicates.

### ACE2-SIT1 expression and purification

Expi293F GnTI− cells in Freestyle 293 Expression Medium (Thermo Fisher Scientific) were transduced with the P3 baculovirus for ACE2 and SIT1 (1.5% v/v for each virus) in the presence of 5 mM sodium butyrate. Cells were grown in a humidity-controlled orbital shaker for 72 h at 30°C with 8% CO_2_ before being harvested by centrifugation at 900 g for 15 min, washed with phosphate-buffered saline, and flash-frozen in liquid nitrogen. Cell pellets were stored at −80°C until further use.

Cell pellets expressing full-length ACE2-SIT1 complex were resuspended in a lysis buffer of 50 mM HEPES pH 7.5, 300 mM NaCl, 1.5% glycol-diosgenin (GDN, Anatrace), and cOmplete EDTA-free Protease Inhibitor Cocktail (Roche). Lysate was loaded on Strep-Tactin XT resin (IBA) gravity flow column, washed with column buffer supplemented with 1 mM ATP and 10 mM MgCl_2_, and eluted in column buffer with 50 mM D-biotin. The eluent was immediately loaded on an anti-FLAG M2 affinity resin gravity flow column, washed with column buffer, and eluted in column buffer supplemented with 0.2 mg/mL FLAG peptide. The protein was further purified by size exclusion chromatography using a Superose 6 Increase (10/300) GL column pre-equilibrated with SEC buffer.

### Cryo-EM sample preparation and data collection

Peak fractions of purified ACE2-SIT1 complex were pooled, incubated with 10 mM L-pipecolate or 10 mM glycine, and concentrated to ∼4.5 mg/mL or ∼6 mg/mL. cryo-EM grids were prepared using a Mark IV Vitrobot (Thermo Fisher Scientific) by applying protein to glow-discharged QuantiFoil Au R1.2/1.3 200-mesh grids (Quantifoil), blotting for 3.0 s under 100% humidity at 4°C, and then plunging into liquid ethane.

The pipecolate dataset was collected on a Titan Krios electron microscope, using a GIF-Quantum energy filter with a 20 eV slit width (Gatan) and a K3 direct electron detector (Gatan) at a dose rate of 19.5e^-^/px/sec. EPU (Thermo Fisher Scientific) was used to automatically record three movie stacks per hole (super-resolution / EPU bin 2) with the defocus ranging from −1.2 to −2.4 µm. Each micrograph was dose-fractioned into 50 frames, with an accumulated dose of 50 e^-^/Å^2^.

The glycine dataset was collected on a Titan Krios electron microscope, using a GIF-Quantum energy filter with a 5eV slit width (Gatan) and a K2 direct electron detector (Gatan). SerialEM was used to automatically record three movie stacks per hole ^62^, at a dose rate of 8.42 e^-^/px/sec with the defocus ranging from −1.0 to −2.2 µm. Each micrograph was dose-fractioned into 50 frames, with an accumulated dose of 50 e^-^/Å^2^.

### Reconstruction of ACE2-SIT1 with pipecolate

cryoSPARC was used for the majority of the data processing workflow ^63^, with RELION used only for final 3D classification (without alignment) for the focused refinement of the SIT1 component ^64^. Movies were motion corrected and CTF-corrected in cryoSPARC.

For the pipecolate dataset, particles were blob picked, followed by two cycles of 2D classification. The well-resolved 2D classes were used for template-based picking. *Ab initio* models generation and heterogeneous classification yielded maps with open and closed conformation of the ACE2 peptidase domain. The particles from each conformation were separated by further classification into species with 2:2 and 2:1 stoichiometries of ACE2 and SIT1. Non-uniform refinement with C2 symmetry imposed gave reconstructions for the 2:2 ACE2:SIT1 open and closed PD conformations at 3.29 Å and 3.24 Å, respectively.

To improve the map for SIT1, all particles from the 2:2 open and closed reconstructions were aligned with C2 symmetry imposed to give a consensus 2:2 ACE2-SIT1 reconstruction. The aligned particles were symmetry-expanded and local refinement was performed within a region encompassing a single SIT1 monomer. The resultant aligned particles were then subjected to 3D classification in RELION without further alignment (K=10, T=12). The particles in the best 3D classes, based on estimated resolution criteria, were pooled for local refinement in cryoSPARC using the SIT1 monomer mask to produce a 3.49 Å reconstruction.

### Reconstruction of ACE2-SIT1 in glycine

For the apo dataset, particles were blob picked, followed by two cycles of 2D classification. The particles from well-resolved 2D classes were used for Topaz training and particle picking ^65^. Subsequent two cycles of *ab initio* model generation and heterogenous refinement yielded maps with open and closed conformation of the ACE2 peptidase domain. The particles from each conformation were further classified into species with 2:2 and 2:1 stoichiometries of ACE2 and SIT1. Non-uniform refinement with C2 symmetry imposed gave reconstructions for the 2:2 ACE2:SIT1 open and closed PD conformations at 3.59 Å and 3.46 Å, respectively.

To improve the map for SIT1, all particles from the 2:2 open and closed reconstructions were aligned with C2 symmetry imposed to give a consensus 2:2 ACE2-SIT1 reconstruction. The aligned particles were symmetry-expanded and local refinement was performed within a region encompassing a single SIT1 monomer. The resultant aligned particles were then subjected to 3D classification in RELION without further alignment (K=10, T=12). The particles in the best 3D classes, based on estimated resolution criteria, were pooled for local refinement in cryoSPARC using the SIT1 monomer mask to produce a 3.76 Å reconstruction.

### Model building and refinement

Models were initially built for the open and closed ACE2 dimer using the pipecolate dataset. Published structures of B^0^AT1 (PDB: 6M18), ACE2 with PD open (PDB: 6M1D), and ACE2 with PD closed (PDB: 6M18) were used as templates for model building. The B^0^AT1-derived SIT1 model was pruned using CHAINSAW ^66^. SIT1 residues 10-582 and ACE2 residues 740-768 were built using the 3.49 Å transmembrane domain-focused map in Coot ^67^. Models were refined with phenix.real_space_refine using default geometric restraints ^68^. For the open and closed 2:2 ACE2-SIT1 complexes, the focused SIT1 coordinates were used as a reference model during refinement. Geometric restraints for pipecolate were generated using GRADE ^69^. The pipecolate-bound ACE2 and SIT1 protein models were used as templates for subsequent model building for ACE2-SIT1 structure determined in the presence of glycine. The ACE2 components required minimal adjustments and differences in the SIT1 were primarily localized around TM1a.

### B^0^AT1-leucine refitting

The B^0^AT1-leucine complex from 6M17 was subjected to global real-space refinement against the deposited focused cryo-EM map (EMD-30043) at 3.1Å using PHENIX. The binding mode of the substrate leucine (Leu707) was then adjusted to maximize its interactions within the binding site while remaining consistent with the cryo-EM density. Alterations were also made to B^0^AT1 substrate pocket which included flipping the peptide backbone of residues Gly51 and Leu52 as well as changes to the sidechain rotamers of Val50, Leu52, Val55 and Trp56. The refitted model was also subjected to global real-space refinement against the deposited focused cryo-EM map. Qscores were calculated for the re-refined and refitted/re-refined B^0^AT1 models using MapQ ^50^.

## Data availability

The cryo-EM maps and models generated in this study have been deposited in the EMDB database and the Protein Data Bank, respectively, under accession codes, EMD-17381 and PDB-8P30 for ACE2-SIT1 with pipecolate bound and open peptidase domain, EMD-17382 and PDB-8P31 for ACE2-SIT1 with pipecolate bound and closed peptidase domain, EMD-17380 and PDB-8P2Z for SIT1 with pipecolate bound focused refinement, EMD-17378 and PDB-8P2X for ACE2-SIT1 with open peptidase domain determined in the presence of glycine, EMD-17379 and PDB-8P2Y for ACE2-SIT1 with closed peptidase domain determined in the presence of glycine, EMD-17377 and PDB-8P2W for SIT1 focused refinement determined in the presence of glycine.

## Acknowledgements

This work was financially supported by the BBSRC (BB/V018051/1) (to E.P.C. and T.D.). D.B.S., J.H., and A.C.W.P. were supported by the Innovative Medicines Initiative 2 Joint Undertaking (JU) under grant agreement No 875510. The JU receives support from the European Union’s Horizon 2020 research and innovation program and EFPIA and Ontario Institute for Cancer Research, Royal Institution for the Advancement of Learning McGill University, Kungliga Tekniska Hoegskolan, Diamond Light Source Limited. We thank Beth Maclean for assistance with EM sample preparation and screening, Loic Carrique for helping with OPIC data collection setup, Brian Marsden for assistance with cryo-EM data processing resources, and Wyatt Yue for assistance with project management. Electron microscopy was provided through the Oxford Particle Imaging Centre (OPIC), an Instruct-ERIC centre (funded by Wellcome Trust JIF award [060208/Z/00/Z] and equipment grant [093305/Z/10/Z]) and the Electron Bio-Imaging Centre, Diamond Light Source Ltd (eBIC; BAG proposal bi28713).

## Author contributions

H.Z.L, J.S.H., S.R.B., D.W., and L.S. cloned SIT1 and ACE2. H.Z.L., J.S.H., S.G.L., K.E.J.R., S.R.B., D.S., A.E., D.H., and C.N. expressed and purified the proteins. H.Z.L. collected and analyzed the thermostabilization data. H.Z.L., G.C., and A.C.W.P. collected and processed the cryo-EM images and built the atomic models. H.Z.L., A.C.W.P., and D.B.S. analyzed the structures. H.Z.L., A.C.W.P., and D.B.S. wrote the manuscript. All authors participated in the discussion and manuscript editing. N.A.B.B., T.R.D., E.P.C., and D.B.S. supervised the research.

## References

1. Richardson, J. S. The Anatomy and Taxonomy of Protein Structure. in Advances in Protein Chemistry vol. 34 167–339 (Elsevier, 1981).

2. Chou, P. Y. & Fasman, G. D. Conformational parameters for amino acids in helical, β-sheet, and random coil regions calculated from proteins. Biochemistry 13, 211–222 (1974).

3. Adzhubei, A. A. & Sternberg, M. J. E. Left-handed Polyproline II Helices Commonly Occur in Globular Proteins. Journal of Molecular Biology 229, 472– 493 (1993).

4. Brodsky, B. & Ramshaw, J. A. M. The collagen triple-helix structure. Matrix Biology 15, 545–554 (1997).

5. Gustafson, C. L. et al. A Slow Conformational Switch in the BMAL1 Transactivation Domain Modulates Circadian Rhythms. Molecular Cell 66, 447–457.e7 (2017).

6. Sarkar, P., Reichman, C., Saleh, T., Birge, R. B. & Kalodimos, C. G. Proline cis-trans Isomerization Controls Autoinhibition of a Signaling Protein. Molecular Cell 25, 413–426 (2007).

7. Fischer, G. & Schmid, F. X. The mechanism of protein folding. Implications of in vitro refolding models for de novo protein folding and translocation in the cell. Biochemistry 29, 2205–2212 (1990).

8. Henzi, V., Reichling, D. B., Helm, S. W. & MacDermott, A. B. L-proline activates glutamate and glycine receptors in cultured rat dorsal horn neurons. Mol Pharmacol 41, 793–801 (1992).

9. Bae, M. et al. SLC6A20 transporter: a novel regulator of brain glycine homeostasis and NMDAR function. EMBO Mol Med 13, (2021).

10. Namavar, Y. et al. Psychiatric phenotypes associated with hyperprolinemia: A systematic review. Am J Med Genet 186, 289–317 (2021).

11. Stevens, B. R. & Wright, E. M. Substrate specificity of the intestinal brush-border proline/sodium (IMINO) transporter. J. Membrain Biol. 87, 27–34 (1985).

12. Kowalczuk, S. et al. Molecular cloning of the mouse IMINO system: an Na+-and Cl−-dependent proline transporter. Biochemical Journal 386, 417–422 (2005).

13. Takanaga, H., Mackenzie, B., Suzuki, Y. & Hediger, M. A. Identification of Mammalian Proline Transporter SIT1 (SLC6A20) with Characteristics of Classical System Imino. Journal of Biological Chemistry 280, 8974–8984 (2005).

14. Bröer, S. et al. Iminoglycinuria and hyperglycinuria are discrete human phenotypes resulting from complex mutations in proline and glycine transporters. J. Clin. Invest. 118, 3881–3892 (2008).

15. Hysi, P. G. et al. Metabolome Genome-Wide Association Study Identifies 74 Novel Genomic Regions Influencing Plasma Metabolites Levels. Metabolites 12, 61 (2022).

16. Yin, X. et al. Genome-wide association studies of metabolites in Finnish men identify disease-relevant loci. Nat Commun 13, 1644 (2022).

17. Kim, J.-H. et al. A Genome-Wide Association Study Identifies Potential Susceptibility Loci for Hirschsprung Disease. PLoS ONE 9, e110292 (2014).

18. Lee, J. S. et al. Association Analysis of SLC6A20 Polymorphisms With Hirschsprung Disease. Journal of Pediatric Gastroenterology & Nutrition 62, 64– 70 (2016).

19. Xie, X. et al. Associations of SLC6A20 genetic polymorphisms with Hirschsprung’s disease in a Southern Chinese population. Bioscience Reports 39, BSR20182290 (2019).

20. Chao, J. R. et al. Human retinal pigment epithelial cells prefer proline as a nutrient and transport metabolic intermediates to the retinal side. Journal of Biological Chemistry 292, 12895–12905 (2017).

21. Strunnikova, N. V. et al. Transcriptome analysis and molecular signature of human retinal pigment epithelium. Human Molecular Genetics 19, 2468–2486 (2010).

22. Bennis, A. et al. Comparison of Mouse and Human Retinal Pigment Epithelium Gene Expression Profiles: Potential Implications for Age-Related Macular Degeneration. PLoS ONE 10, e0141597 (2015).

23. Bonelli, R. et al. Identification of genetic factors influencing metabolic dysregulation and retinal support for MacTel, a retinal disorder. Commun Biol 4, 274 (2021).

24. Gao, X. R., Huang, H. & Kim, H. Genome-wide association analyses identify 139 loci associated with macular thickness in the UK Biobank cohort. Human Molecular Genetics 28, 1162–1172 (2019).

25. Vuille-dit-Bille, R. N. et al. Human intestine luminal ACE2 and amino acid transporter expression increased by ACE-inhibitors. Amino Acids 47, 693–705 (2015).

26. Shen, Y. et al. Structures of ACE2–SIT1 recognized by Omicron variants of SARS-CoV-2. Cell Discov 8, 123 (2022).

27. The Severe Covid-19 GWAS Group. Genomewide Association Study of Severe Covid-19 with Respiratory Failure. N Engl J Med 383, 1522–1534 (2020).

28. COVID-19 Host Genetics Initiative et al. Mapping the human genetic architecture of COVID-19. Nature 600, 472–477 (2021).

29. The GenOMICC Investigators et al. Genetic mechanisms of critical illness in COVID-19. Nature 591, 92–98 (2021).

30. Bröer, S. & Gether, U. The solute carrier 6 family of transporters: The Solute Carrier Family 6. British Journal of Pharmacology 167, 256–278 (2012).

31. Yamashita, A., Singh, S. K., Kawate, T., Jin, Y. & Gouaux, E. Crystal structure of a bacterial homologue of Na+/Cl--dependent neurotransmitter transporters. Nature 437, 215–223 (2005).

32. Malinauskaite, L. et al. A mechanism for intracellular release of Na+ by neurotransmitter/sodium symporters. Nat Struct Mol Biol 21, 1006–1012 (2014).

33. Gotfryd, K. et al. X-ray structure of LeuT in an inward-facing occluded conformation reveals mechanism of substrate release. Nat Commun 11, 1005 (2020).

34. Krishnamurthy, H. & Gouaux, E. X-ray structures of LeuT in substrate-free outward-open and apo inward-open states. Nature 481, 469–474 (2012).

35. Piscitelli, C. L. & Gouaux, E. Insights into transport mechanism from LeuT engineered to transport tryptophan: Insights into transport mechanism. The EMBO Journal 31, 228–235 (2012).

36. Focht, D. et al. A non-helical region in transmembrane helix 6 of hydrophobic amino acid transporter MhsT mediates substrate recognition. EMBO J 40, (2021).

37. Weyand, S. et al. Structure and Molecular Mechanism of a Nucleobase–Cation– Symport-1 Family Transporter. Science 322, 709–713 (2008).

38. Faham, S. et al. The Crystal Structure of a Sodium Galactose Transporter Reveals Mechanistic Insights into Na ^+^ /Sugar Symport. Science 321, 810–814 (2008).

39. Yan, R. et al. Structural basis for the recognition of SARS-CoV-2 by full-length human ACE2. Science 367, 1444–1448 (2020).

40. Yan, R. et al. Structural basis for the different states of the spike protein of SARS-CoV-2 in complex with ACE2. Cell Res 31, 717–719 (2021).

41. Chen, Y. et al. ACE2-targeting monoclonal antibody as potent and broad-spectrum coronavirus blocker. Sig Transduct Target Ther 6, 315 (2021).

42. Motiwala, Z. et al. Structural basis of GABA reuptake inhibition. Nature 606, 820– 826 (2022).

43. Coleman, J. A., Green, E. M. & Gouaux, E. X-ray structures and mechanism of the human serotonin transporter. Nature 532, 334–339 (2016).

44. Yang, D. & Gouaux, E. Illumination of serotonin transporter mechanism and role of the allosteric site. Sci. Adv. 7, eabl3857 (2021).

45. Munck, B. G. Transport of imino acids and non-α-amino acids across the brush-border membrane of the rabbit ileum. J. Membrain Biol. 83, 15–24 (1985).

46. Olkhova, E., Raba, M., Bracher, S., Hilger, D. & Jung, H. Homology Model of the Na+/Proline Transporter PutP of Escherichia coli and Its Functional Implications. Journal of Molecular Biology 406, 59–74 (2011).

47. Bröer, A. et al. Sodium translocation by the iminoglycinuria associated imino transporter (SLC6A20). Molecular Membrane Biology 26, 333–346 (2009).

48. Shajahan, A. et al. Comprehensive characterization of N-and O-glycosylation of SARS-CoV-2 human receptor angiotensin converting enzyme 2. Glycobiology 31, 410–424 (2021).

49. D’Imprima, E. et al. Protein denaturation at the air-water interface and how to prevent it. eLife 8, e42747 (2019).

50. Pintilie, G. et al. Measurement of atom resolvability in cryo-EM maps with Q-scores. Nat Methods 17, 328–334 (2020).

51. Kantcheva, A. K. et al. Chloride binding site of neurotransmitter sodium symporters. Proc. Natl. Acad. Sci. U.S.A. 110, 8489–8494 (2013).

52. Sauer, D. B. et al. Structural basis of ion – substrate coupling in the Na+-dependent dicarboxylate transporter VcINDY. Nat Commun 13, 2644 (2022).

53. Wang, X. & Boudker, O. Large domain movements through the lipid bilayer mediate substrate release and inhibition of glutamate transporters. eLife 9, e58417 (2020).

54. Sauer, D. B. et al. Structure and inhibition mechanism of the human citrate transporter NaCT. Nature 591, 157–161 (2021).

55. Nayal, M. & Cera, E. D. Valence Screening of Water in Protein Crystals Reveals Potential Na+Binding Sites. Journal of Molecular Biology 256, 228–234 (1996).

56. Zhao, Y. et al. Single-molecule dynamics of gating in a neurotransmitter transporter homologue. Nature 465, 188–193 (2010).

57. Shafqat, S. et al. Human brain-specific L-proline transporter: molecular cloning, functional expression, and chromosomal localization of the gene in human and mouse genomes. Mol Pharmacol 48, 219–229 (1995).

58. Pei, J., Kim, B.-H. & Grishin, N. V. PROMALS3D: a tool for multiple protein sequence and structure alignments. Nucleic Acids Research 36, 2295–2300 (2008).

59. Price, M. N., Dehal, P. S. & Arkin, A. P. FastTree 2 – Approximately Maximum-Likelihood Trees for Large Alignments. PLoS ONE 5, e9490 (2010).

60. Chi, G. et al. Phospho-regulation, nucleotide binding and ion access control in potassium-chloride cotransporters. EMBO J 40, (2021).

61. Mahajan, P., et al. Expression Screening of Human Integral Membrane Proteins Using BacMam. in Structural Genomics (eds. Chen, Y. W. & Yiu, C.-P. B.) vol. 2199 95–115 (Springer US, 2021).

62. Schorb, M., Haberbosch, I., Hagen, W. J. H., Schwab, Y. & Mastronarde, D. N. Software tools for automated transmission electron microscopy. Nat Methods 16, 471–477 (2019).

63. Punjani, A., Rubinstein, J. L., Fleet, D. J. & Brubaker, M. A. cryoSPARC: algorithms for rapid unsupervised cryo-EM structure determination. Nat Methods 14, 290–296 (2017).

64. Zivanov, J., Nakane, T. & Scheres, S. H. W. Estimation of high-order aberrations and anisotropic magnification from cryo-EM data sets in RELION -3.1. IUCrJ 7, 253–267 (2020).

65. Bepler, T. et al. Positive-unlabeled convolutional neural networks for particle picking in cryo-electron micrographs. Nat Methods 16, 1153–1160 (2019).

66. Stein, N. CHAINSAW : a program for mutating pdb files used as templates in molecular replacement. J Appl Crystallogr 41, 641–643 (2008).

67. Emsley, P. & Cowtan, K. Coot : model-building tools for molecular graphics. Acta Crystallogr D Biol Crystallogr 60, 2126–2132 (2004).

68. Afonine, P. V. et al. Real-space refinement in PHENIX for cryo-EM and crystallography. Acta Crystallogr D Struct Biol 74, 531–544 (2018).

69. Smart, O. S. et al. Grade. https://www.globalphasing.com (2011).

