## Supplementary figures for "Structure and function of the SIT1 proline transporter in complex with the COVID-19 receptor ACE2"

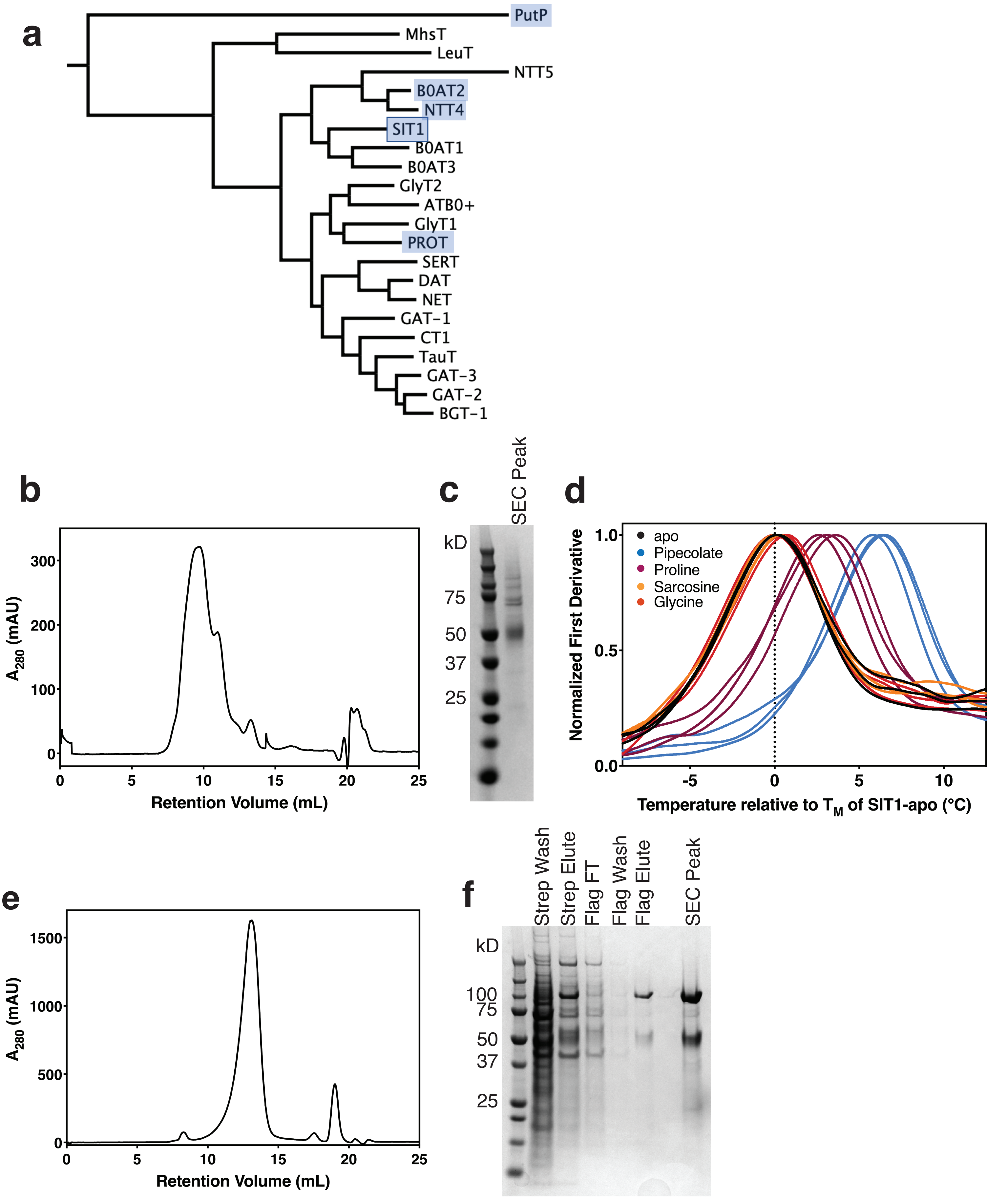


**Supplementary Figure 1. Purification of SIT1 and the ACE2-SIT1 complex.** (**a**) Phylogenetic tree of human SLC6 proteins and bacterial homologs, with proline transporters highlighted. (**b**) Preparative size-exclusion chromatography of SIT1 on a Superdex 200 Increase 10/300 GL column. (**c**) Representative SDS-PAGE of SIT1 purification from 10 biological replicates. (**d**) Thermostabilization of detergent solubilized SIT1 by various amino acids. (**e**) Preparative size-exclusion chromatography of the ACE2-SIT1 complex on a Superose 6 Increase 10/300 GL column. (**f**) Representative SDS-PAGE for the purification of the ACE2-SIT1 complex from 10 biological replicates.

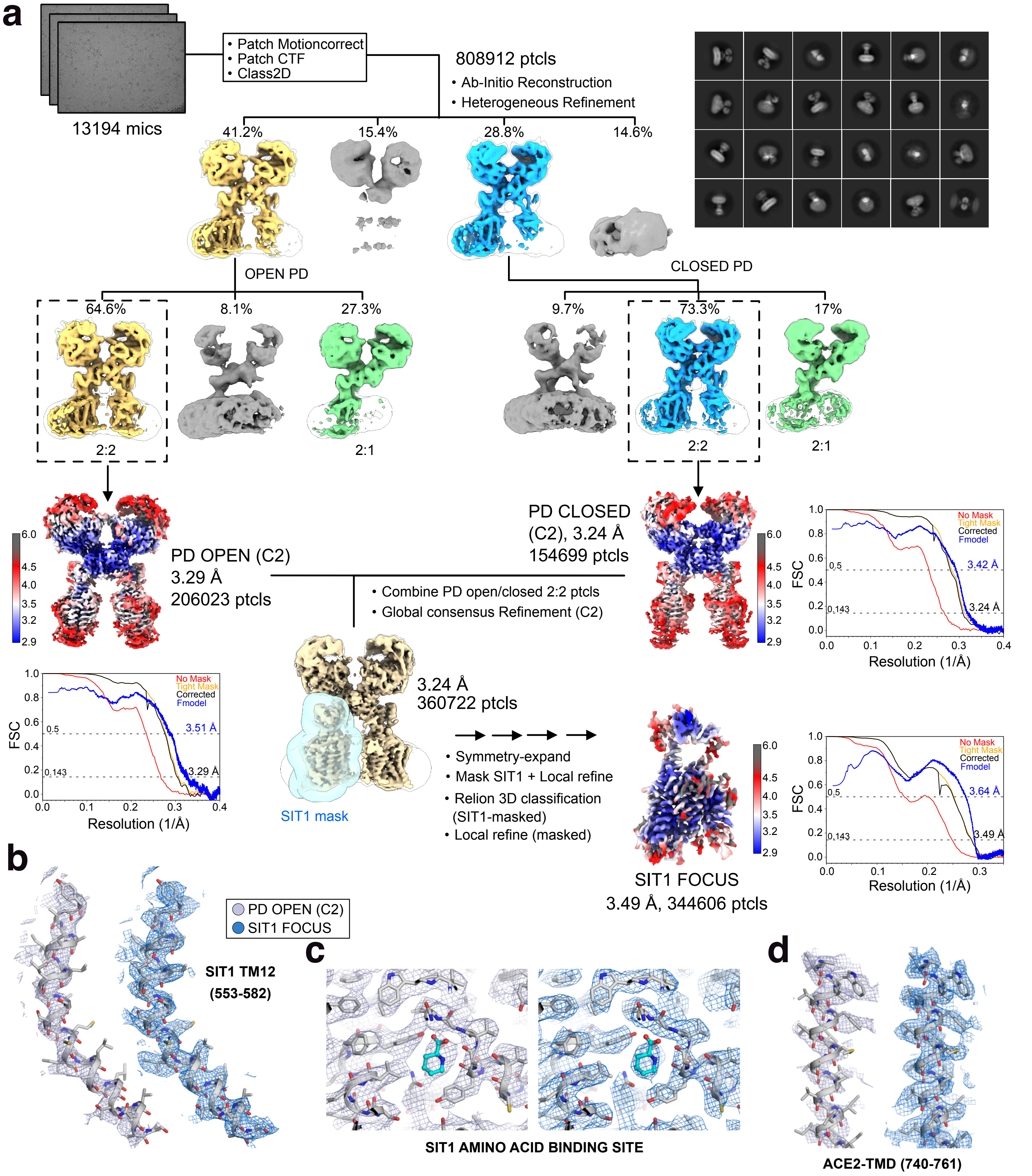


**Supplementary Figure 2. ACE2-SIT1 structure determination.** (**a**) Flow chart for determining the structure of the ACE2-SIT1 complex in the presence of pipecolate. Representative densities for (**b**) SIT1's TM12, (**c**) SIT1's amino acid binding site, and (**d**) ACE2's TMD. Coulombic potential maps are shown in grey and blue respectively for the PD open map with C2 symmetry imposed and the SIT1 focused refinement, respectively.

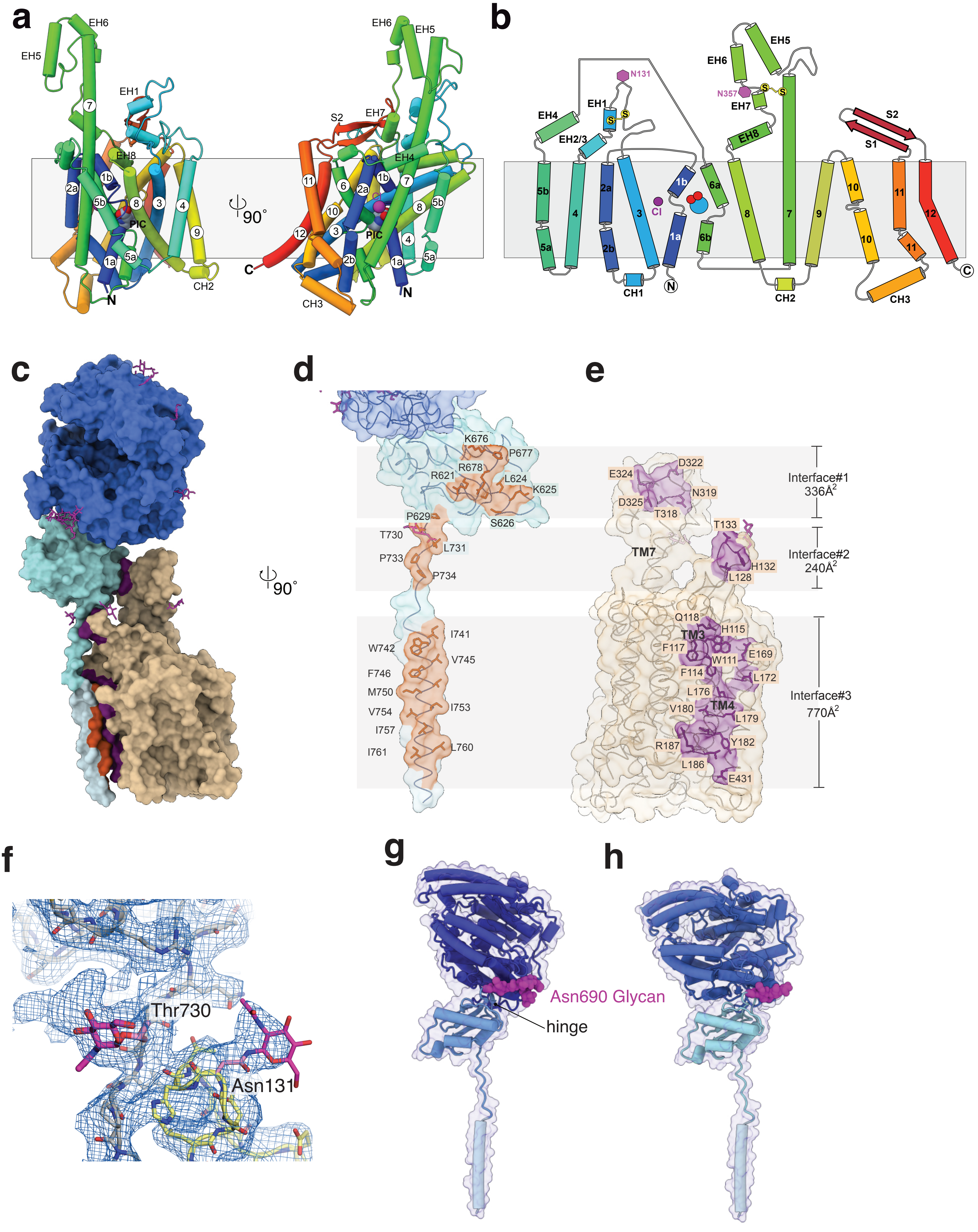


**Supplementary Figure 3. ACE2-SIT1 interactions.** SIT1's (**a**) heterodimeric structure and (**b**) topology. (**c**) ACE2-SIT1 dimeric structure. ACE2 and SIT1 colored in blues and wheat, respectively. ACE2 and SIT1 interface residues colored in orange and purple, respectively. Open book representation for dimer interface residues of (**d**) ACE2 and (**e**) SIT1. (**f**) O-linked glycosylation at Thr730. ACE2 structure, SIT1 structure, and Coulombic potential map colored in grey, yellow, and blue, respectively. ACE2 structure viewed from the plane of the membrane (**g**) open and (**h**) closed conformation. Glycan at Asn690 shown as magenta spheres.

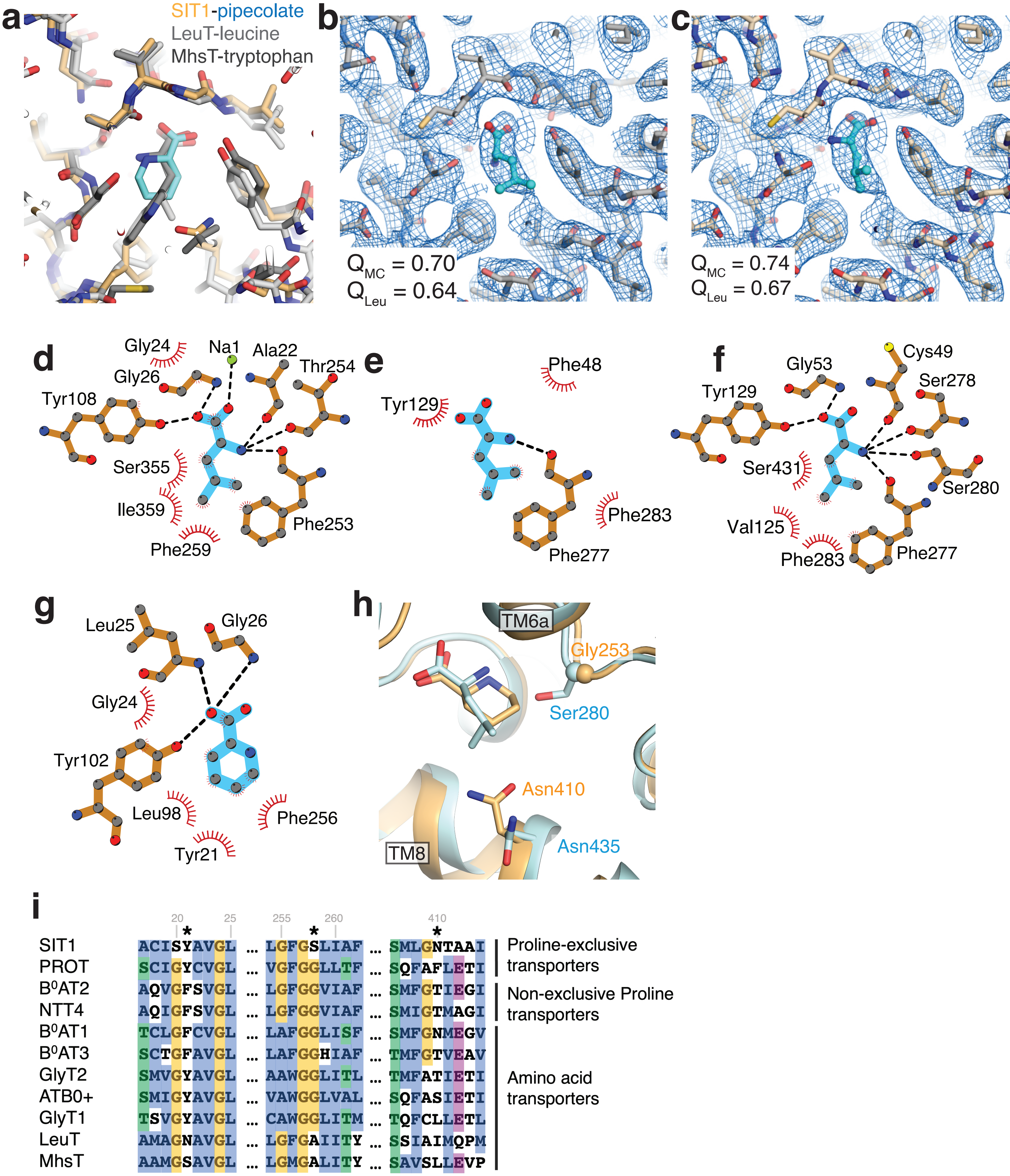


**Supplementary Figure 4. SLC6 amino acid interactions.** (**a**) Overlay of amino-acid bound structures of SIT1, LeuT (2A65), and MhsT (4US4). (**b**) Published structure of leucine bound B$^0$AT1 (6M17) with map-to-model fit scores for the main chain and leucine substrate. (**c**) Refit structure of leucine bound B$^0$AT1 with map-to-model fit scores for the main chain and leucine substrate. Contact maps for (**d**) leucine-bound LeuT, (**e**) published leucine-bound B$^0$AT1 (6M17), (**f**) refit leucine-bound B$^0$AT1, and (**g**) pipecolate bound SIT1. (**h**) Overlay of pipecolate bound SIT1 and leucine-bound B$^0$AT1, colored in wheat and blue, respectively. (**i**) Sequence alignment for SLC6 amino acid transporters and homologs.

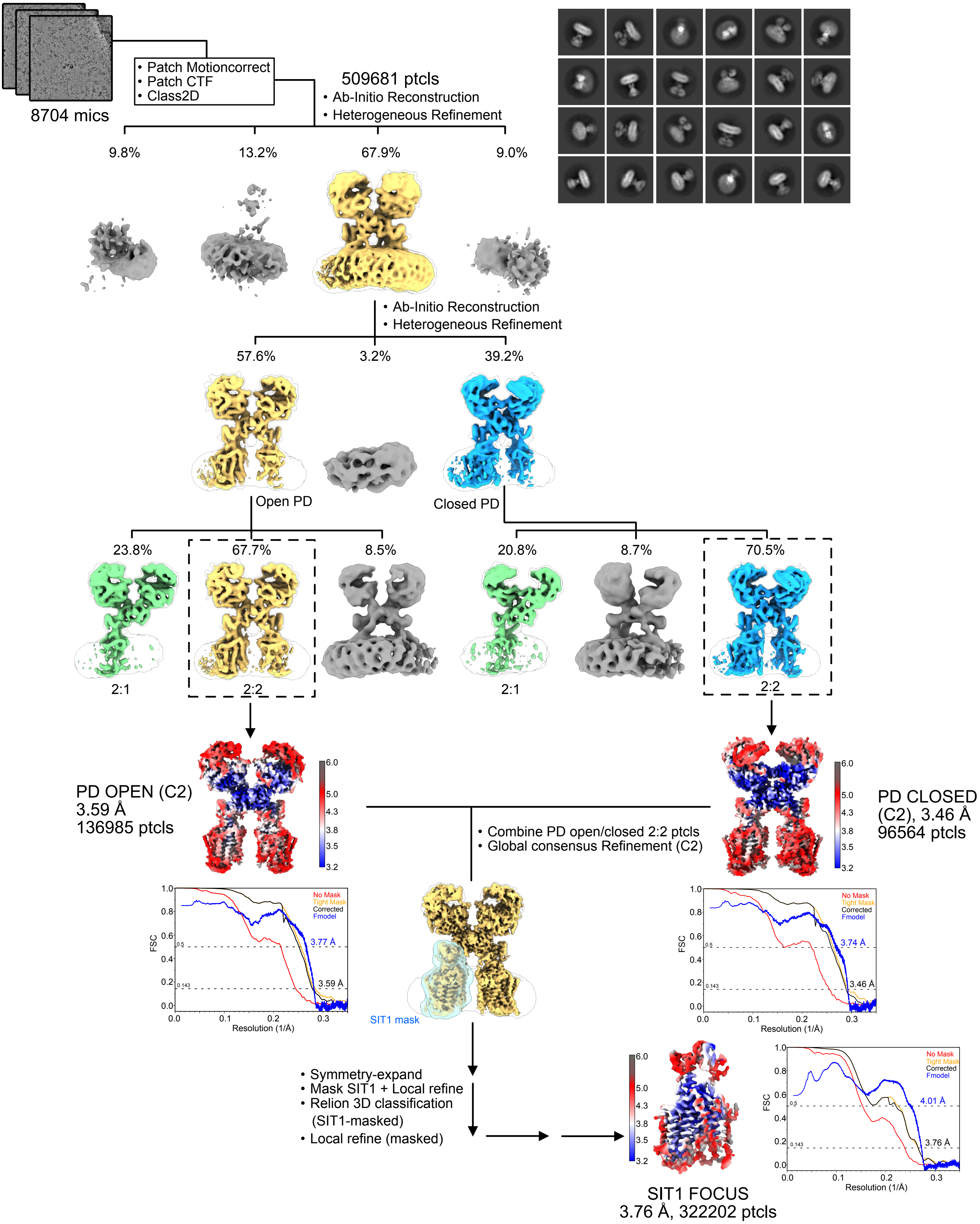


**Supplementary Figure 5. ACE2-SIT1 apo structure determination.** Flow chart for determining the structure of the ACE2-SIT1 complex in the presence of glycine.

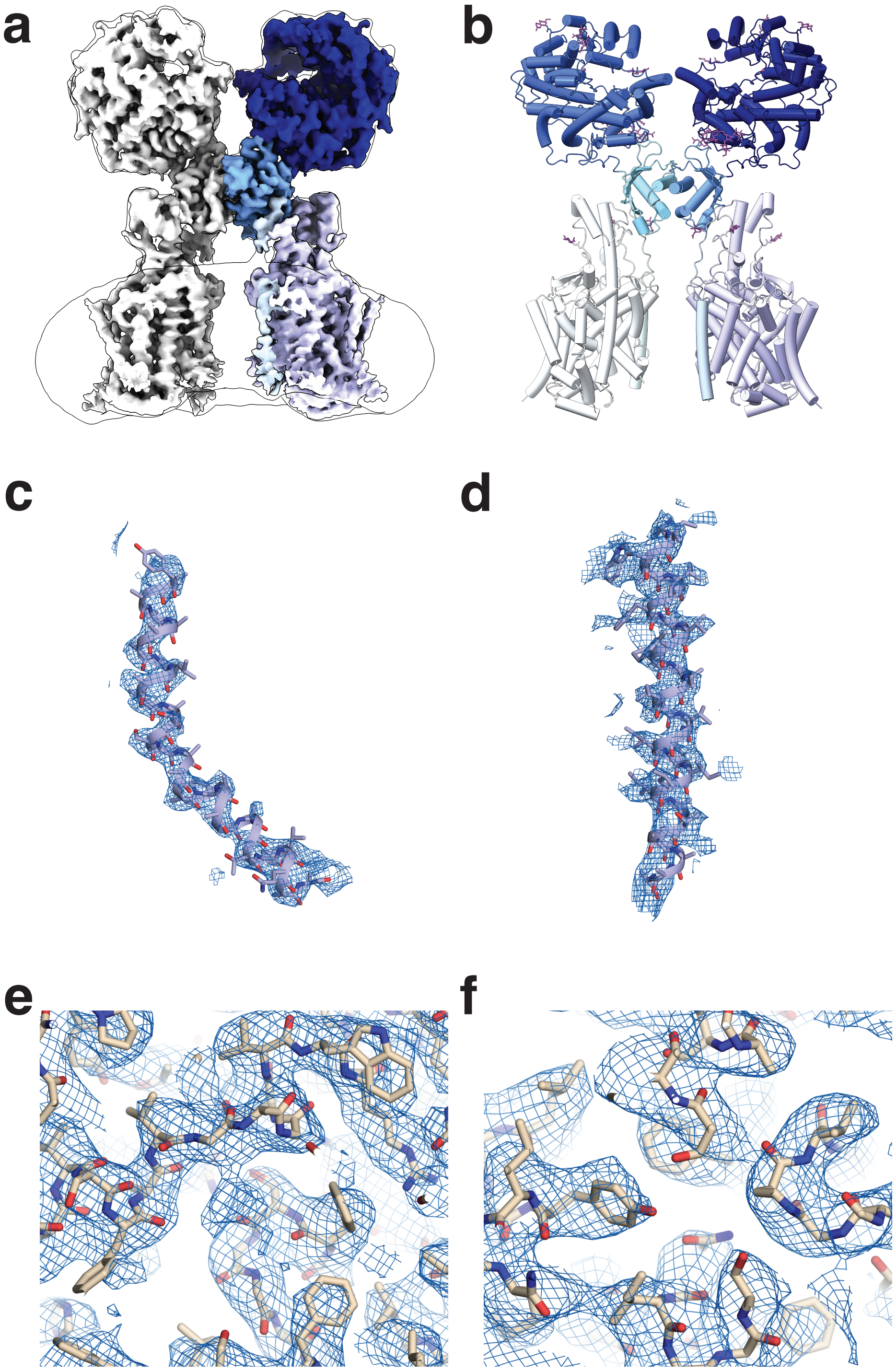


**Supplementary Figure 6. SIT1-apo structure and binding sites.** (**a**) Coulombic potential map and (**b**) experimental model for ACE2-SIT1 determined in the presence of glycine. Representative densities for (**c**) SIT1's TM12 and (**d**) ACE2's TMD. (**e**) Amino acid and (**f**) chloride binding sites of SIT1, determined in the presence of glycine. Coulombic potential map shown as blue mesh, protein shown as sticks.
